# Optimal microbial pathway variants can be determined by large-scale bioenergetic evaluation in syntrophic propionate oxidation

**DOI:** 10.1101/2020.02.07.939629

**Authors:** Mauricio Patón, Héctor H. Hernández, Jorge Rodríguez

## Abstract

The complete understanding of microbial propionate oxidation in syntrophy with hydrogenotrophic methanogenesis remains elusive due to uncertainties in pathways and mechanisms for interspecies electron transfer (IET). Possible pathway variants differ in their intermediate metabolites, on which electron carriers are involved and in which steps are coupled to (and to how many) proton translocations. In this work, a systematic methodology was developed (based on sound biochemical, physiological and bioenergetic principles) to evaluate the feasibility and net ATP yield of large sets of pathway variants under different physiological and environmental conditions. A pathway variant is deemed feasible under given conditions only if all pathway reaction steps have non-positive Gibbs energy change and if all the metabolite concentrations remain within an acceptable physiological range (10^−6^ to 10^−2^ M). Several million combinations of pathway variants and parameters/conditions were evaluated for propionate oxidation, providing an unprecedented mechanistic insight into its biochemical and bioenergetic landscape. Propionate oxidation via lactate appeared as the most ATP yielding pathway under most of the conditions evaluated. Results under typical methanogenic conditions indicate that syntrophic propionate oxidation can sustain life only at hydrogen partial pressures within the range of 1.2 to 4 Pa. These extremely low concentrations constitute a kinetic impossibility and strongly suggest for IET mechanisms other than dissolved hydrogen.

**Importance:** In this work an original methodology was developed that quantifies the bioenergetically and physiologically feasible net ATP yields for large numbers of microbial metabolic pathways and their variants under different conditions. This ensures global optimality in finding the pathway variant(s) leading to the highest ATP yield. The methodology is especially relevant to hypothesise which microbial pathway variants are most likely to prevail in microbial ecosystems under high selective pressure for efficient metabolic energy conservation.

Syntrophic microbial oxidation of propionate to acetate has extremely low energy available and requires very high metabolic efficiency in order to sustain life. Our results bring mechanistic insights into the optimum pathway variants and the impact of environmental conditions on the ATP yields and other metabolic bottlenecks. Additionally, our results conclude that IET mechanisms other than hydrogen must exist to simultaneously sustain the growth of both propionate oxidisers and hydrogenotrophic methanogens.

## Introduction

Propionate oxidation to acetate and hydrogen is a highly endergonic reaction under standard conditions (ΔG^01^ = +76.1 kJ/mol propionate (1)). The reaction however, can become exergonic and yield sufficient energy for net ATP production only at very low hydrogen partial pressures (P_H2_) (1-10 Pa) typical in methanogenic environments (2–4). In the case of the reduction reaction of CO_2_ to methane via hydrogen, although it is highly exergonic under standard conditions (ΔG^01^= −131 kJ/mol; (1)), under typical methanogenic conditions (very low P_H2_) the reaction falls much closer to equilibrium with actual energy available between −15 and −40 kJ/mol (3). Both volatile fatty acid (VFA) oxidisers and methanogens are known to grow very close to thermodynamic equilibrium (5).

Due to these bioenergetic limitations, propionate oxidation is believed to be possible only under syntrophic association with hydrogen scavenging microorganisms (6). The specialised nature of methanogenic archaea, which are able to grow only on very few substrates (acetate, CO_2_, hydrogen/formate or other C1 compounds such as methanol) (4,7), makes them dependent on other microorganisms for their supply of substrate. Both syntrophic reactions can proceed simultaneously only within a narrow range of concentrations if dissolved hydrogen is the interspecies electron transfer (IET) mechanism, this range is known as the methanogenic niche. The fact that, under methanogenic conditions, the amount of energy available from either of the two syntrophic reactions is smaller than the minimum needed for one ATP unit synthesis via substrate level phosphorylation, implies that metabolic energy conservation must be driven by chemiosmotic transmembrane proton translocations (1).

Significant work has been done on the elucidation of electron transfer mechanisms between syntrophic partners; propionate (or butyrate) oxidisers with methanogens. Different mechanisms for IET have been proposed to occur via hydrogen and/or formate. Although IET via hydrogen has been identified as more suitable than formate due to its higher diffusivity (8), IET via formate has also been proposed when microorganisms do not grow in aggregates, given its much higher solubility (6,9–12). Formate and hydrogen production in the same microorganism have also been proposed to take place at different reaction sites with (i) formate produced at the reoxidation step of menaquinone from the oxidation of succinate to fumarate and (ii) hydrogen produced in the reoxidation of the NADH from the malate oxidation to oxaloacetate and the ferredoxin reduction of pyruvate to acetyl-CoA reactions, respectively (4,13). The hypothesis of both IET capable species simultaneously produced is supported by faster observed growth in presence of syntrophic methanogens that metabolize both hydrogen and formate (14,15). Formate has been suggested to serve in those cases as a temporary electron sink (16). Alternative mechanisms of IET have been proposed including via conductive pili (also called nanowires) as in *Geobacter sulfurreducens* (17–19). *Geobacter* species appears to be the only bacteria known to date to display this feature, which appears to facilitate the microbial growth in aggregates (20). *Methanobacterium* (hydrogen or formate utilising methanogens) appears to be in low abundance when microorganisms growth in aggregates with *Geobacter* species, suggesting that IET should occur via an alternative mechanism to H_2_ or formate (21). Evidence of syntrophic partners transferring electrons via shuttle molecules (sulfur species, humic substances or flavins), through nanowires or conductive minerals have also been reported (22–24). Methanogenesis rates from propionate and acetate have shown to increase significantly after addition of iron oxides during growth (25).

Numerous studies have focused on elucidating the catabolic pathways of propionate oxidation to acetate and numerous different possible pathways have been described including: (i) propionate oxidation via the methylmalonyl-CoA pathway, which has been extensively studied (4,6,10,11,13,16,26–30), (ii) propionate oxidation via lactate (26,31,32) or (iii) propionate oxidation via hydroxypropionyl-CoA (26,31). Propionate oxidisers that use the methylmalonyl-CoA pathway are however the only ones that have been isolated (6). The conversion of propionate via an alternative butyrate and acetate yielding pathway has also been reported (33,34).

Although detailed thermodynamic studies have been conducted of individual reactions present in related microbial catabolic pathways (35–40), the complete understanding of many microbial conversions remains unachieved. This is largely due to the lack of clarity on the different possible pathway variants and/or mechanisms that drive endergonic reactions. Pathway variants are defined here in terms of which intermediate metabolites, in particular electron carriers, are involved as well as in terms of the mechanisms for energy conservation (i.e. proton translocations) and the location in which they take place within a pathway. In order to address these remaining uncertainties in terms of pathway variants and syntrophic IET, a comprehensive bioenergetic evaluation of a very large set of reported pathway variants is presented in this work for propionate oxidation as well as for hydrogenotrophic methanogenesis. The consideration of the impact that intermediate metabolite concentrations have on the bioenergetics of each reaction step across each pathway is central to determine the feasibility of each pathway variant and the quantification of its net ATP yield. The syntrophic pathways evaluation for an ample range of hydrogen partial pressures is specifically targeted to understand the limits of the IET mechanism and of the methanogenic niche within which syntrophic propionate oxidisers and methanogens can both simultaneously sustain growth.

## Methodology

### Selection of pathways for propionate oxidation

The selection of the catabolic pathways to be considered for the oxidation of propionate to acetate as per Eq. 1 was compiled through comprehensive literature review (4,6,11,13,26,28,31).

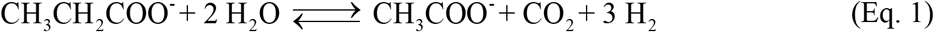

Pathways yielding stoichiometries different to that of Eq. 1, such as propionate conversion via butyrate into acetate (33,34) were not included in this work.

Literature has been found reporting variability in terms of which electron carriers are involved in certain reaction steps of the methylmalonyl-CoA pathway. In the oxidation of succinate to fumarate, menaquinone has been reported as the electron carrier (11,16) while FADH_2_ has also been reported as possible electron carrier for the same reaction step (27). Discrepancies in the specific terminal products from electron carrier reoxidation were also found. The oxidation of NADH carriers has been proposed to occur through hydrogenases (16,41–43). Formate dehydrogenases have also been reported to oxidise menaquinone (11,30). Hydrogen and formate however, appear to be thermodynamically equivalent (13,44) and therefore only hydrogen was considered in this work as the terminal product of the electron carrier oxidations.

In the case of hydrogenotrophic methanogenesis, the pathway was also compiled from literature (2,3,7,45–47) and, specifically, the energy conservation sites via proton translocation (48,49).

Selected pathway reactions were cross referenced from the literature sources and the Kyoto Encyclopaedia of Genes and Genomes (KEGG) database (50). Only reactions based on enzymes reported in prokaryotes were considered. Although for some cases microorganisms known to carry out an entire pathway may have not been yet isolated (e.g. P_5_ or P_6_ in Table 1), in this work, all reactions in a given pathway variant were assumed to take place within a single cell. It is worth noting that some of the pathways selected such as methylmalonyl-CoA (P_2_,P_3_), lactate (P_4_) or hydroxypropionyl-CoA (P_6_) contain cyclic steps (4,16,28). The complete set of pathways considered for the oxidation of propionate are presented in Table 1 and those for hydrogenotrophic methanogenesis in Table 2. Graphical representations of the pathways are available in the Supplementary Information Figures S1.1-1.6.

**Table 1.**
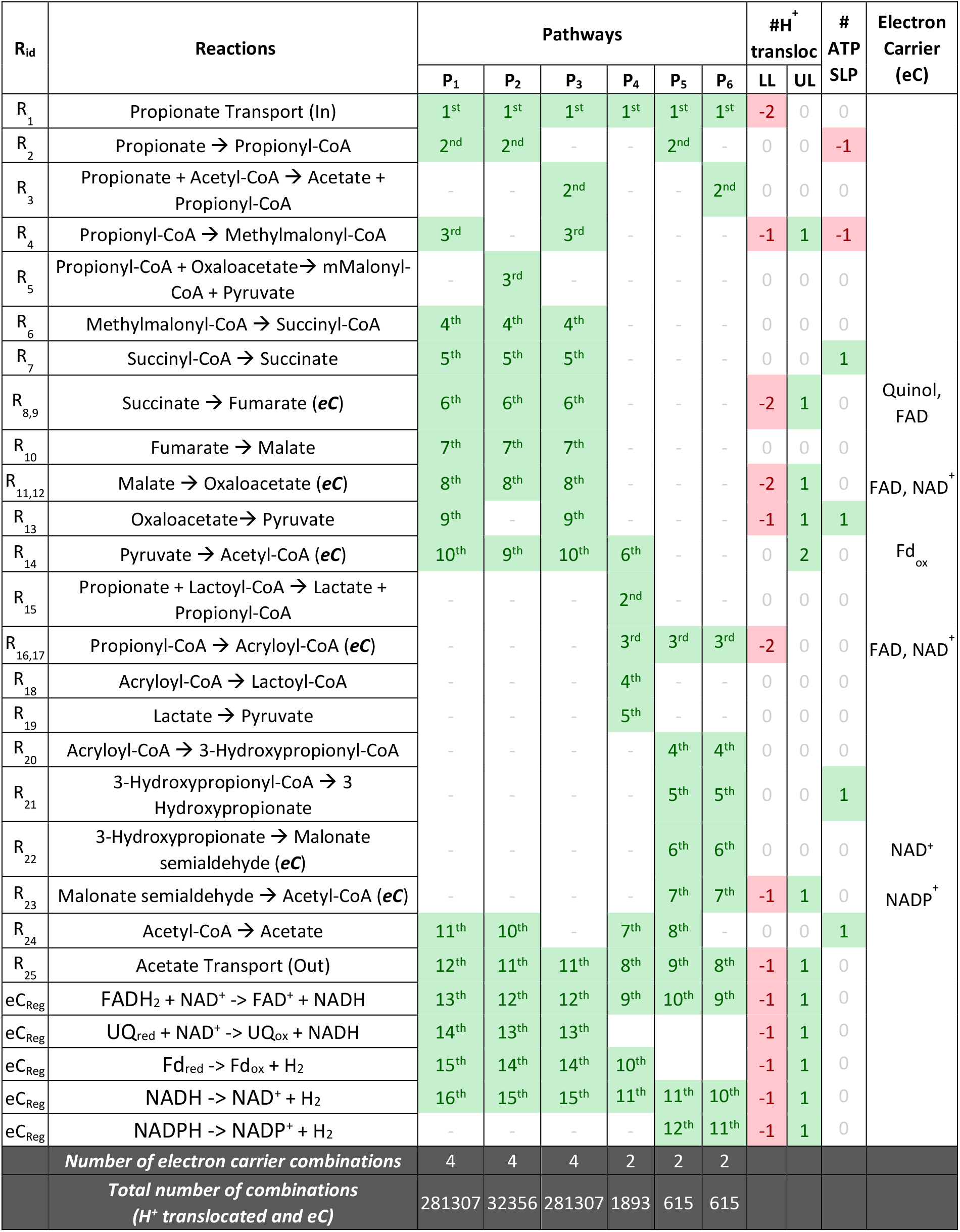
The complete set of pathway reactions considered for propionate oxidation to acetate as per Eq. 1 are presented. The reactions considered for electron carrier regeneration are also included (eC_Reg_). The numbers under the pathways (P_n_) indicate the order in which reactions occur in each pathway. Lower (LL) and upper limits (UL) for the number of protons translocations in a specific reaction step, ATP consumption/production as substrate level phosphorylation (SLP) for each reaction are indicated.

**Table 2.**
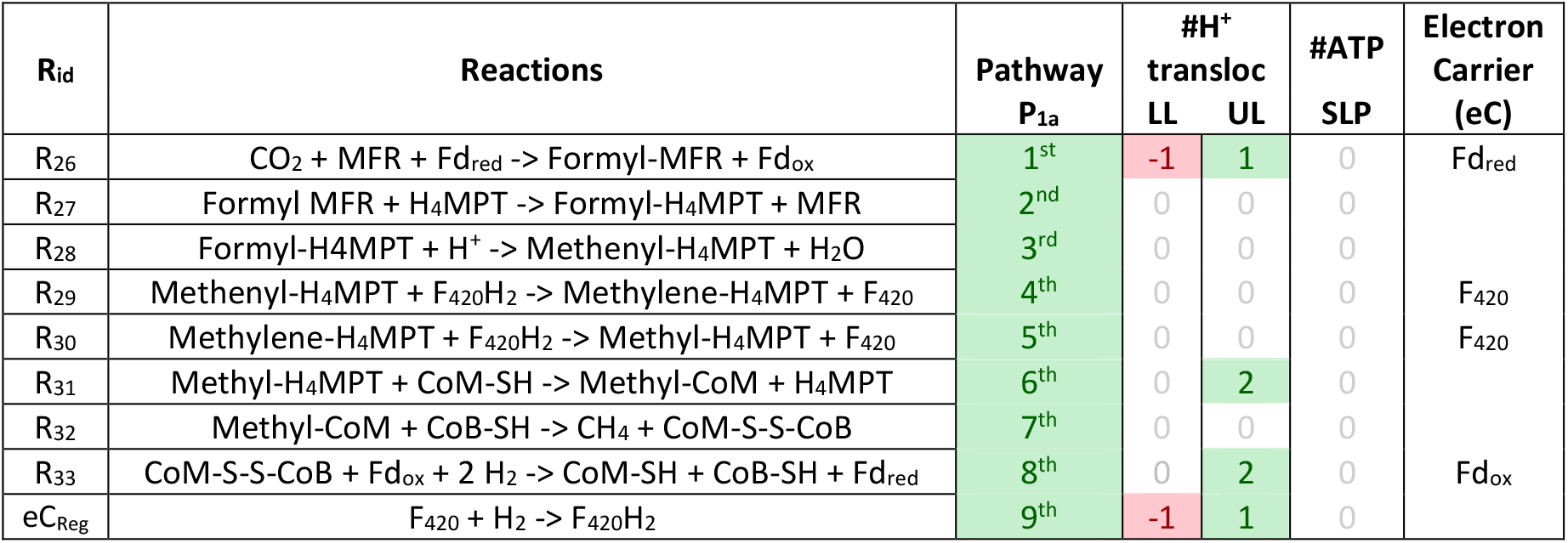
The set of reactions considered for CO_2_ reduction with H_2_ to methane are presented. The reactions considered for electron carrier regeneration are also included (eC_Reg_). The numbers under the pathways (P_n_) indicate the order in which reactions occur in each pathway. Lower (LL) and upper limits (UL) for the number of protons translocations is a specific step are indicated. No ATP consumption/production via substrate level phosphorylation (SLP) was reported for this reaction.

### Environmental conditions for pathway evaluation

A set of environmental conditions representative of a typical stable methanogenic anaerobic digestion steady state operation were selected as reference (51). The pathway variants are evaluated at those constant conditions and extracellular concentrations of substrates and products. The propionate and acetate concentration values selected were set at 1.4·10^−4^ M and 3.07·10^−3^ M respectively. The hydrogen partial pressure (P_H2_) was set to a default 1.62 Pa and is assumed to be in equilibrium with its corresponding dissolved concentration.

### Intracellular metabolite concentrations

Based on the values for intracellular metabolite concentrations reported in literature (52) and on theoretical calculations (53), all internal metabolite concentrations were constrained within a physiologically feasible maximum of 10^−2^ M and a minimum of 10^−6^ M. The small volume of a cell (circa 1 μm^3^) (54) implies that less than one hundred single molecules would be present inside the cell at 10^−7^ M a number considered too low for any feasible subsequent positive reaction rate. The total concentrations of other conserved moieties such as electron carriers and free CoA were defined as parameters (38,55). The concentrations of electron carrier were characterised by the ratios between their reduced and oxidised forms and constrained such that their total concentration is conserved and neither form falls outside the physiological limits described above (this implies maximum and minimum reduced/oxidised carrier ratios of 10^−4^ and 10^4^ respectively).

### Thermodynamic parameters and assumptions

The thermodynamic values of Gibbs free energy (and enthalpies) of formation, required for the thermodynamic calculations for each reaction step, were collected from the literature for each metabolite (1,3,56–59). The enthalpy of formation values of a limited number of metabolites involved in the oxidation of propionate to acetate were unavailable and had to be estimated. Detailed references for the thermodynamic parameters along with the estimation methods used for some enthalpies, are provided in the Supplementary Information S3. Temperature corrected bioenergetics were applied to all pathways reactions for propionate oxidation using the Van’t Hoff equation. Temperature corrections were however not applicable to the hydrogenotrophic methanogenesis pathway reactions, due to the unavailability of enthalpies for methanofuran (MFR) or tetrahydromethanopterin (H_4_MPT), which are important in pathway.

### Chemiosmotic energy conservation

All reactions identified to take place via membrane-bound enzymes are assumed, in principle, as capable of proton translocation through the cell membrane to, either directly recover energy as proton motive force (pmf), or to drive endergonic reactions in a pathway. Those energy conservation sites were identified both through previous literature (1,4,48,60–65) and the online database Metacyc (66).

### Assessment of pathways feasibility

For each reaction step in which an electron carrier was involved, a set of possible electron carriers variants were defined. Additionally, for each reaction step with proton translocation capability, a range of possible numbers of proton translocations that can take place in that step were defined (see Table 1).

All possible variants, combinatorial of all electron carrier variations with all possible numbers of proton translocation in the capable steps, were evaluated for each pathway. The feasibility of any given pathway variant aligns with minimum energy dissipation and catabolism efficiency and is therefore evaluated by seeking a zero or minimum Gibbs energy dissipation in all pathway reactions steps. This criterion allows for the sequential calculation of the corresponding product concentrations of each pathway step given the substrate from the previous step.

The evaluation of a pathway variant consisted first of the determination of its feasibility in terms of whether for all reaction steps to have a zero or negative Gibbs energy change, all metabolite intermediate concentrations can still remain within the defined physiological limits. A pathway variant is deemed unfeasible and is discarded if any of the metabolite concentrations must fall below the lower physiological limit in order to thermodynamically enable a preceding reaction to occur. If a reaction step has high energy and allows (maintaining still ΔG<0) for the produced metabolite to take concentration values higher than the upper physiological limit (10^−2^ M), the concentration sits at the upper limit and energy is dissipated and lost. The evaluation of a pathway variant that is feasible concludes with the quantification of its overall net ATP yield. A pseudo code representation of the algorithm developed is shown in the Supplementary Information S3.1.

Some of the pathways evaluated contain cycles (e.g. P_2_-P_4_ from Table 1). A pathway contains a cycle when one reaction in the pathway requires two substrates to yield two products (excluding from here the conserved moieties such as electron carriers and free CoA). A specific section of the algorithm had to be developed to be able to evaluate the cyclic steps and metabolite concentrations based exactly on the same principles described above and not involving additional assumptions (see Supplementary Information S3.1).

In addition to cycles, pathways can contain electron bifurcation reactions (65,67) as is the case in the reduction of the CoM-CoB heterodisulfide in the methanogenesis pathway, (R_34_, Table 2). This allows for the reduction of CO_2_ to Formyl-MFR via the produced reduced ferredoxin (R_26_, Table 2). A specific section of the algorithm was also developed in order to evaluate pathways in which electron bifurcation takes place (see Supplementary Information S3.2).

The combinatory set of possible pathway variants as defined above becomes very large (nearly 600 000 in this case) for each set of physiological parameters and environmental conditions as defined in Table 3. The automation capacities of the algorithm developed allowed for the evaluation of the complete domain of all possible pathway variants. This ensures global optimality since the pathway variants with the highest ATP yields found have to be indeed the optima in terms of metabolic energy conservation.

**Table 3.** Set of physiological parameters and environmental conditions evaluated for propionate oxidisers and hydrogenotrophic methanogens. The values indicated were evaluated for each parameter individually leaving all other parameters at the default reference value (bold font and grey shaded). Label (a) refers to default values for propionate oxidisers and label (b) refers to default values for hydrogenotrophic methanogens. CoA-SH and CoM-SH refer to the concentrations of their free forms.

|  | Parameters | Units | Values |  |  |  |  |  |  |
| --- | --- | --- | --- | --- | --- | --- | --- | --- | --- |
| Physiological<br>Parameters | $\Delta G_{ATP}$ | kJ/mol | -40 | -45 | **-50<sup>(a)(b)</sup>** | -55 | -60 | -65 | |
|  | H<sup>+</sup>/ATP | [] | **9/3<sup>(b)</sup>** | **10/3<sup>(a)</sup>** | 11/3 | 12/3 | 13/3 | 14/3 | 15/3 |
|  | CoA-SH | mM | 10<sup>-3</sup> | 10<sup>-2</sup> | 10<sup>-1</sup> | **1<sup>(a)</sup>** | 10 |  |  |
|  | CoM-SH | mM | 10<sup>-3</sup> | 10<sup>-2</sup> | 10<sup>-1</sup> | **1<sup>(b)</sup>** | 10 |  |  |
|  | H<sub>4</sub>MPT | mM | 10<sup>-3</sup> | 10<sup>-2</sup> | 10<sup>-1</sup> | **1<sup>(b)</sup>** | 10 |  |  |
|  | pH<sub>in</sub> | [] | 6 | 6.5 | **7<sup>(a)(b)</sup>** | 7.5 | 8 |  |  |
| Environmental<br>Conditions | H<sub>2</sub> | Pa | 0.01 | **0.13<sup>(a)</sup>** | 1.62 | **31.63<sup>(b)</sup>** | 316.3 |  |  |
|  | CO<sub>2</sub> | mM | 10<sup>-3</sup> | 10<sup>-2</sup> | 10<sup>-1</sup> | 1 | **10<sup>(a)(b)</sup>** |  |  |
|  | T<sup>(1)</sup> | °C | **25<sup>(b)</sup>** | **35<sup>(a)</sup>** | 45 | 55 |  |  |  |
|  | pH<sub>out</sub> | [] | 6 | **7<sup>(a)(b)</sup>** | 8 |  |  |  |  |

<sup>(1)</sup>For hydrogenotrophic methanogenesis, a sensitivity analysis was not performed for temperature. All ATP yields are computed with enthalpies at 25 °C

### Parameter selection and sensitivity analysis

The values reported in literature for some of the required physiological parameters show variability (Table 3). Different values of ΔG_ATP_ hydrolysis under physiological conditions have been reported that range from as small as −45 or −50 kJ/mol (1) to −60 to −70 kJ/mol (4). The number of protons translocated per turn of the ATP synthase has been widely reported as 9 protons per turn resulting in 3 ATPs (which leads to the widely accepted ratio of 3 protons per ATP). However, the number of protons required for a complete turn of the ATP synthase is known to vary based on the number of c-subunits able to translocate protons in the ATP synthase (68). This number has been reported to vary from 8 to 15 c-subunits (4,68–72) or equivalent to a ratio of 2.7–5 H^+^/ATP (73). A similar modelling approach was proposed, for a total number of protons per mol of ATP (35). Under their approach, the number of protons per mol of ATP was defined as an integer number. In our approach, a fractional number of protons per ATP is proposed, based in the previously explained total number of protons required for a full turn on the ATP synthase, which results in the generation of 3 ATP. It is worth noting that for a ratio of 15 protons per 3 ATPs and with a ΔG_ATP_ of −50 kJ/mol, the minimum quantum for metabolic energy conservation could be as low as −10 kJ/mol, in line with previous reported values for minimum energy required for microbial growth (5,74). Such low energy quanta could enable energy conservation in microorganisms growing on substrates that yield very low metabolic energy such as propionate.

Intracellular free coenzyme A (CoA-SH) concentrations have been previously reported as high as 10 mM (35) and measured in a butyrate culture to vary between 100-200 μM (75). Due to these differences in values, the impact of the CoA-SH (for propionate oxidisers) and CoM-SH (for methanogens) on the net ATP yields of all pathways was specifically evaluated at different concentrations. Additionally, specific environmental variables such as temperature and pH with possible impact on the bioenergetics were also evaluated.

An overview of the sets of parameter sets evaluated for this sensitivity analysis purpose is shown in Table 3. A total of 30 parameter set configurations were evaluated for all the pathway variants (which corresponded to 18.5 million pathway variant-parameter set scenarios). Within this evaluation space, only from the feasible pathway variants (i.e. all reactions with a ΔG_R_ <= 0 plus all metabolites within the defined physiological limits) and, among these, only those with positive net ATP yield are presented and discussed. All other pathway variants are deemed either unfeasible or unable to sustain microbial growth under the given conditions.

## Results and discussion

The pathways variants with highest ATP yield along with their corresponding metabolite concentration profiles and proton translocation configurations are presented in this section.

### Propionate oxidation

Figure 1 presents the results in terms of net ATP yield of the propionate oxidation pathways evaluated as function of changes in physiological parameters and environmental conditions around the default values in Table 3.

**Figure 1.**
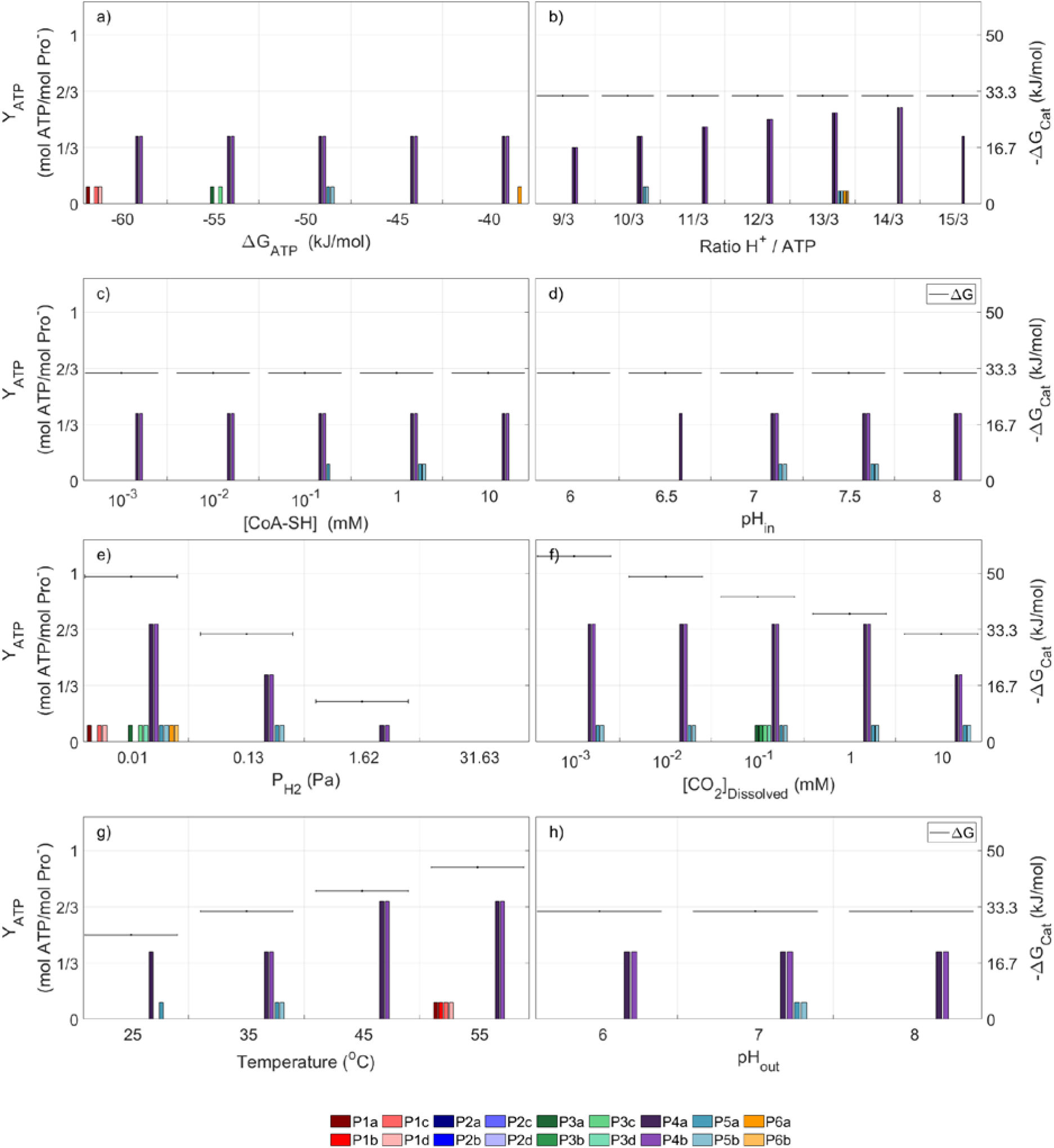
The net ATP produced for each pathway under different physiological (a-d) and environmental (e-h) parameters are shown above. Only the parameter indicated below each graph is modified in respect to the reference conditions (shaded in grey in Table 3). Horizontal lines show the total catabolic energy available (−ΔG_cat_) for pathway efficiency visualisation.

The results consistently present the lactate pathway (P_4a_) as the one biochemically and thermodynamically capable of yielding the most ATP during propionate oxidation. Only if the ΔG_ATP_ were to take values more negative than −45 kJ/mol (Figure 1a), pathways such as that via methylmalonyl-CoA (P_1_,P_3_) or via hydroxypropionyl-CoA (P_5_) appear to yield positive, although smaller, net ATP. This is explained by peering into their intermediate metabolite profiles (see Supplementary Information, Figure S5.3), ΔG_ATP_ values more positive than −55 kJ/mol do not appear to lead to sufficient energy to drive some of the endergonic reactions of the methylmalonyl-CoA pathway (e.g. propionyl-CoA to methylmalonyl-CoA, succinate to fumarate or malate to oxaloacetate).

The value of the ratio H^+^/ATP (Figure 1b) shows that a small energy quantum (up to an optimum ratio H^+^/ATP of 14/3) could extremely favour the efficiency of the lactate pathway with an almost complete conservation of the entire catabolic energy (see Supplementary Information S5). The concentration values of free CoA (Figure 1c) do not appear to impact the efficiency of the lactate pathway in the range of values evaluated. Intracellular pH (Figure 1d) shows that no propionate oxidation pathway variant can possibly produce net ATP at the lowest pH value evaluated (pH 6). The net ATP yields by the lactate pathway appear to be unaffected by pH at values above 6.5. This could be explained by the number of reactions in which net protons are produced in the lactate pathway (R_14_, R_19_, R_24_) (see Supplementary Information, Figure S5.3).

The different environmental conditions considered (see Table 3) imply differences in the overall catabolic energy available. The only exception is perhaps the extracellular pH (Figure 1h) with almost no impact due to the very similar acidity (pK_a_ values) for propionate and acetate (substrate and product of the overall reaction). Lower values of the P_H2_ (Figure 1e), make the overall reaction more exergonic, and potentially more net ATP can be produced. However even at very low P_H2_ values (1.62 Pa), such as found in methanogenic environments, the energy available is below the energy of one net proton translocation. Only a few of the pathway variants that involve the propionyl-CoA to acryloyl-CoA reaction steps (P_4a_, P_5a_) appear as even feasible.

Analogously, for the dissolved external CO_2_ concentration (also a product of the overall reaction) the lower its concentration, the more catabolic energy is available. However, the methylmalonyl-CoA pathway appears to have a bottleneck as it becomes unfeasible below 0.1 mM CO_2_ concentrations. This is explained by the role of CO_2_ in the carboxylation step of propionyl-CoA to methylmalonyl-CoA. If the concentration of CO_2_ is insufficient, the reaction becomes endergonic and the pathway cannot proceed (see Supplementary Information, Figure S5.3). A last impact is that of higher temperature (Figure 1g) which, due to the reaction entropy increase, makes the overall reaction more exergonic, allowing for more pathway variants to reach net ATP yields.

The pathway evaluation method developed provides the metabolite concentrations profiles of all feasible reactions. In Figure 2 the profile is shown for the pathway variants that obtained the highest net ATP yield, namely propionate oxidation via lactate (P_4a_), at three different partial pressures of hydrogen.

**Figure 2.**
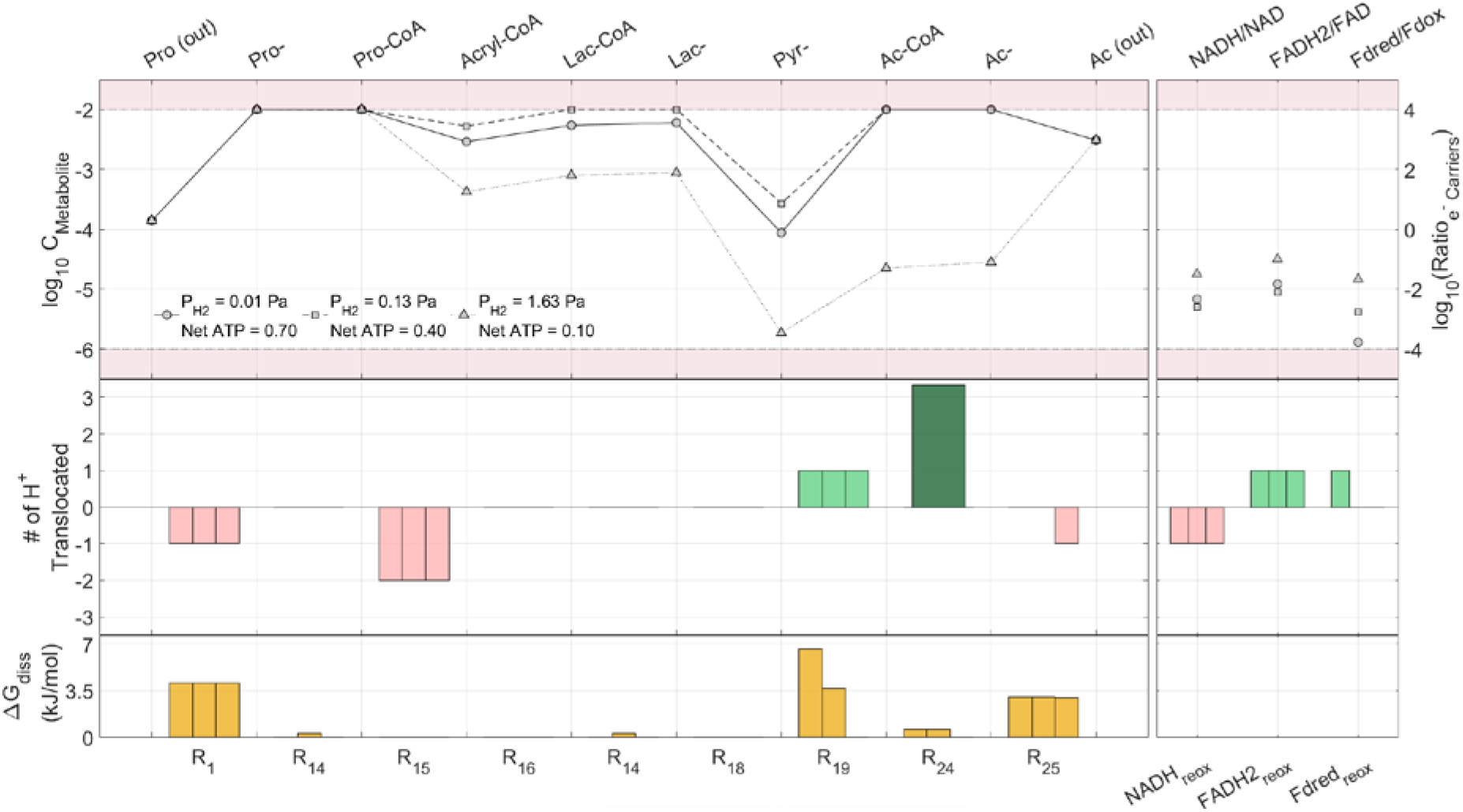
Pathway metabolite concentrations in the propionate oxidation via lactate pathway (P4a) at different hydrogen partial pressures. Symbols in grey (top figure) indicate the logarithmic concentration of the metabolite as labelled in the upper axis. Concentration range outside the physiological limits is the shaded red area. Green and red bars (middle figure) indicate energy conservation reactions in which either energy is recovered or consumed to fuel a reaction, via proton translocations. Darker green bar indicates a reaction in which ATP is produced by substrate level phosphorylation. Yellow bars (bottom figure) indicate Gibbs free energy dissipation (loss) at each reaction steps in the pathway. The physiological parameters and environmental conditions (other than for P_H2_) set as a reference in Table 3 were used.

Figure 2 shows that all metabolites remain within physiological limits for all the P_H2_ values evaluated (same P_H2_ as shown in Figure 1e). As the catabolic energy decreases with increasing product concentration (H_2_), less net energy in the form of translocated protons can be recovered by the cell, particularly in the oxidation of reduced ferredoxin and in the possible transport of acetate outside of the cell. Energy must be and is dissipated (as described in the Methodology section) in those reactions whose products reach the maximum physiological concentrations (e.g. pyruvate to acetyl-CoA at P_H2_ of 1.63 and 0.13 Pa respectively). Figure 2 also clearly illustrates the energetic bottlenecks in the lactate pathway namely (i) the oxidation of propionyl-CoA to acryloyl-CoA, which requires the input of two protons and (ii) the conversion of lactate to pyruvate, which requires an intracellular concentration of pyruvate of already near 10^−6^ M to carry out this reaction at a H_2_ partial pressure of 1.63 Pa.

### Hydrogenotrophic methanogenesis

Figure 3 presents the results in terms of net ATP yield of the evaluated hydrogenotrophic methanogenesis pathway as function of changes in physiological parameters and environmental conditions around the default values in Table 3.

**Figure 3.**
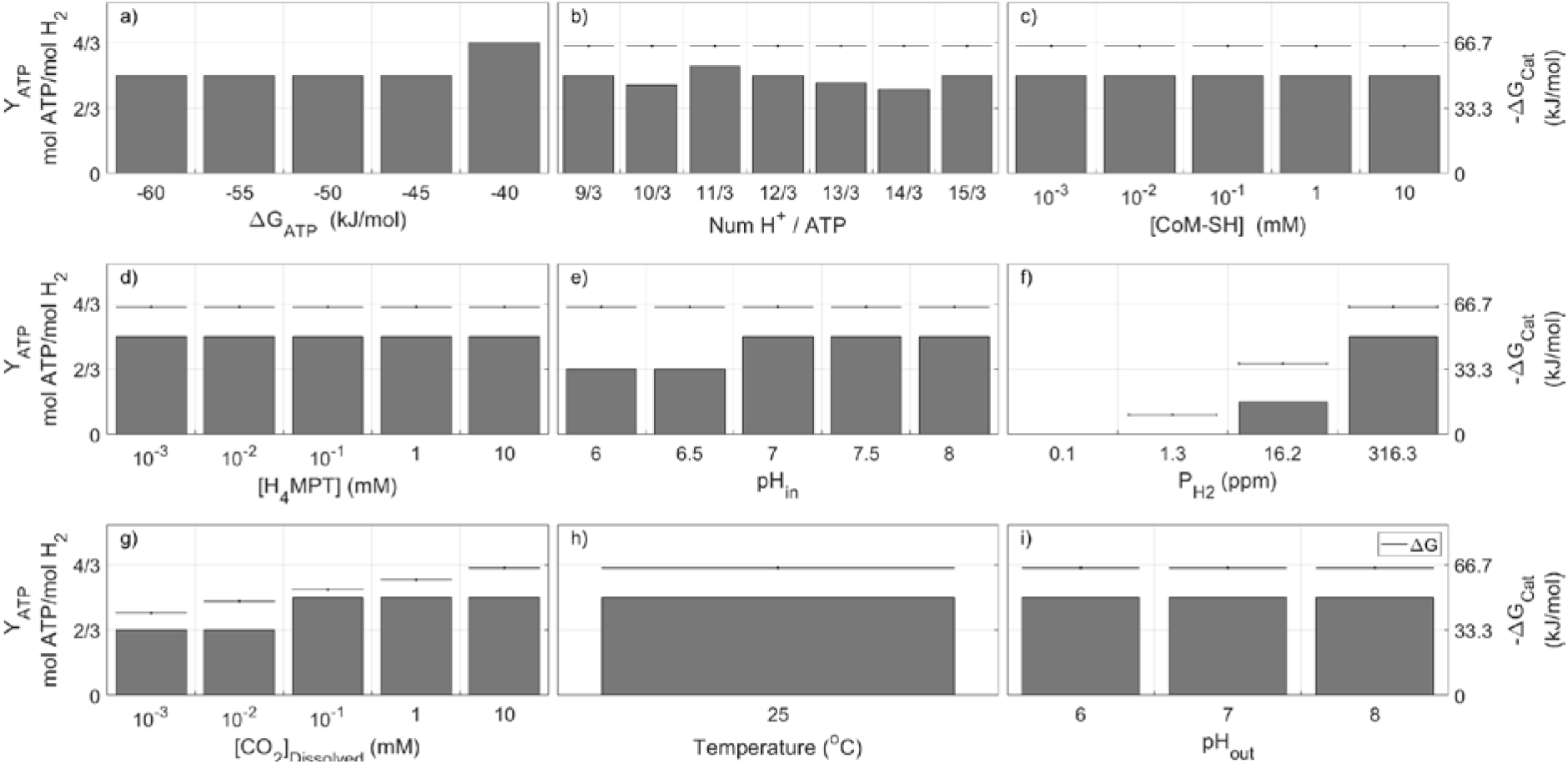
Net ATP equivalents produced in the hydrogenotrophic methanogenesis pathway for different physiological parameters (a-e) and environmental conditions (f-i). In each plot only one parameter, as indicated, is modified respect to the reference conditions from Table 3 (shaded in grey). Temperature (h) could only be evaluated at 25 °C due to the unavailability of enthalpies of formation for certain key compounds involved in the pathway.

As shown in Figure 3, unlike propionate oxidisers and due to the fact that no ATP is produced at substrate level phosphorylation, the value of ΔG_ATP_ only impacts the size of the energy quantum considered in reactions with proton translocation. Hydrogenotrophic methanogenesis displayed no sensitivity to ΔG_ATP_ in the range from −45 to −60 kJ/mol. A quantum of energy of −ΔG_ATP_ of 40 kJ/mol or smaller appears to allow for one additional net proton translocation. In this pathway the optimum H^+^/ATP ratio appears to be 11/3, higher than the ratio of 14/3 for propionate oxidisers. Intracellular pH values lower than 7 appears to decrease the net ATP, while this appears to not be affected by the concentrations of CoM nor of H_4_MPT within the evaluated range of values for those conserved moieties.

As for the case of propionate oxidation, different environmental conditions imply differences in the overall catabolic energy available, except for extracellular pH since no net acids are produced or consumed. Since hydrogen and CO_2_ (Figure 3f and Figure 3g) are substrates of the hydrogenotrophic methanogenesis reaction, the higher their concentration, the higher the catabolic energy available and the higher is potentially the net ATP recovered.

The intracellular metabolite concentration profiles for the hydrogenotrophic methanogenesis pathway are shown in Figure 4 for three values of the hydrogen partial pressures.

**Figure 4.**
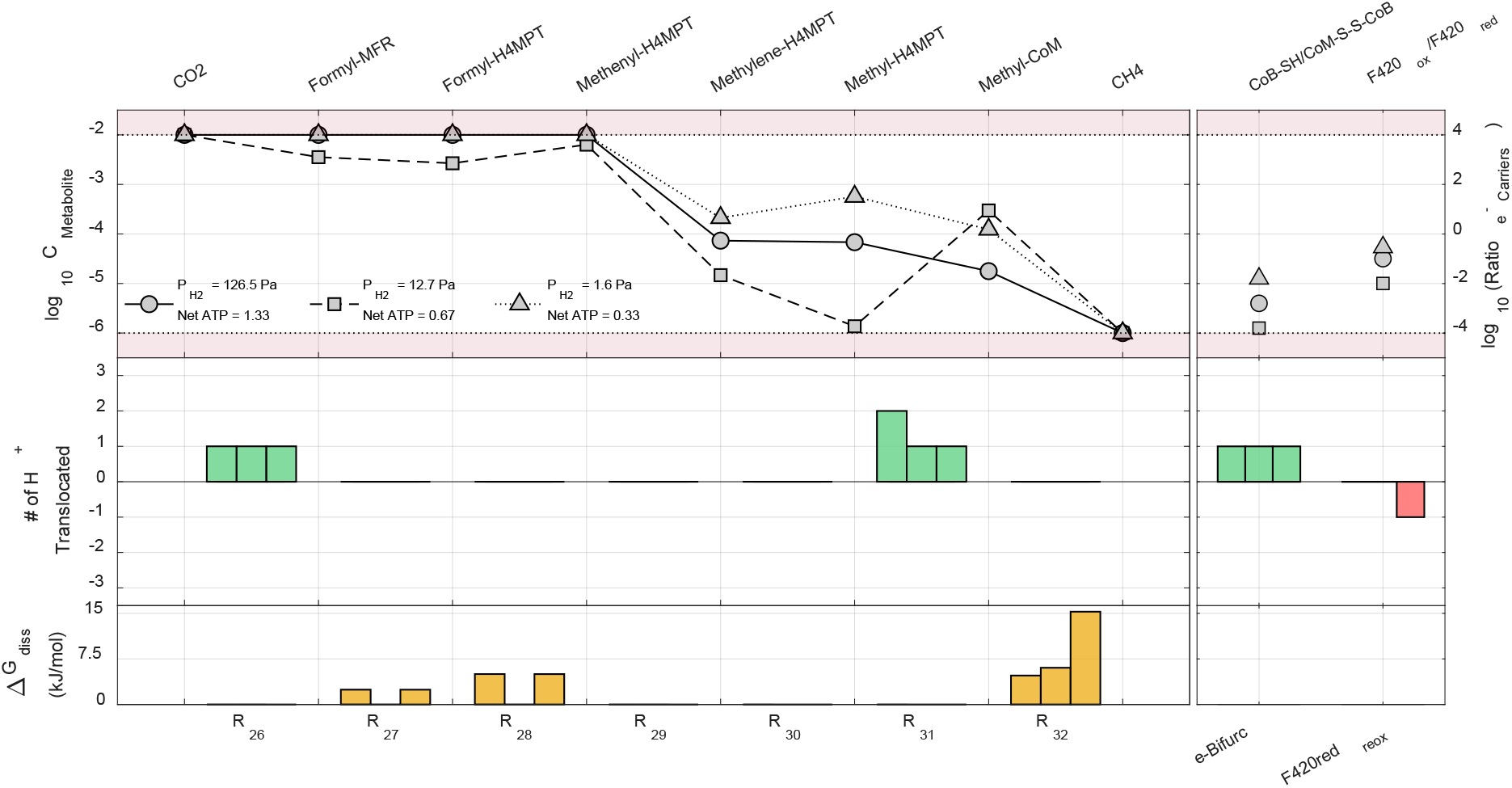
Pathway metabolite concentrations in the hydrogenotrophic methanogenesis pathway. Symbols in grey (top figure) indicate the logarithmic concentration of the metabolite as labelled in the upper axis. Concentration range outside the physiological limits is the shaded red area. Green and red bars (middle figure) indicate energy conservation reactions in which either energy is recovered or consumed to fuel a reaction, via proton translocations. Yellow bars (bottom figure) indicate Gibbs free energy dissipation (loss) at each reaction steps in the pathway. The physiological parameters and environmental conditions (other than for P_H2_) set as a reference in Table 3 were used.

The feasibility and potential bottlenecks in the hydrogenotrophic methanogenesis pathway are shown in Figure 4 along with the sites of energy conservation (via proton translocation). The steps highly impacted by the values of P_H2_ are the reduction of the coenzyme F_420_ (F420red_reox_) and the electron bifurcation reaction of the conversion of CO_2_ to formyl-MFR coupled to the reduction of the CoM-S-S-CoB to CoM-SH (R_26_).

### Syntrophic propionate oxidation and methanogenesis: methanogenic niche

To evaluate the simultaneous syntrophic growth of microorganisms conducting propionate oxidation and hydrogenotrophic methanogenesis, the achievable net ATP yield for each of the two microbial groups was evaluated (Figure 5) as function of the hydrogen concentration (considered the sole mechanism for IET). The reference physiological parameters as per in Table 3 were used for both microbial groups. The typical environmental conditions from a methanogenic anaerobic digestion scenario were used (51).

**Figure 5.**
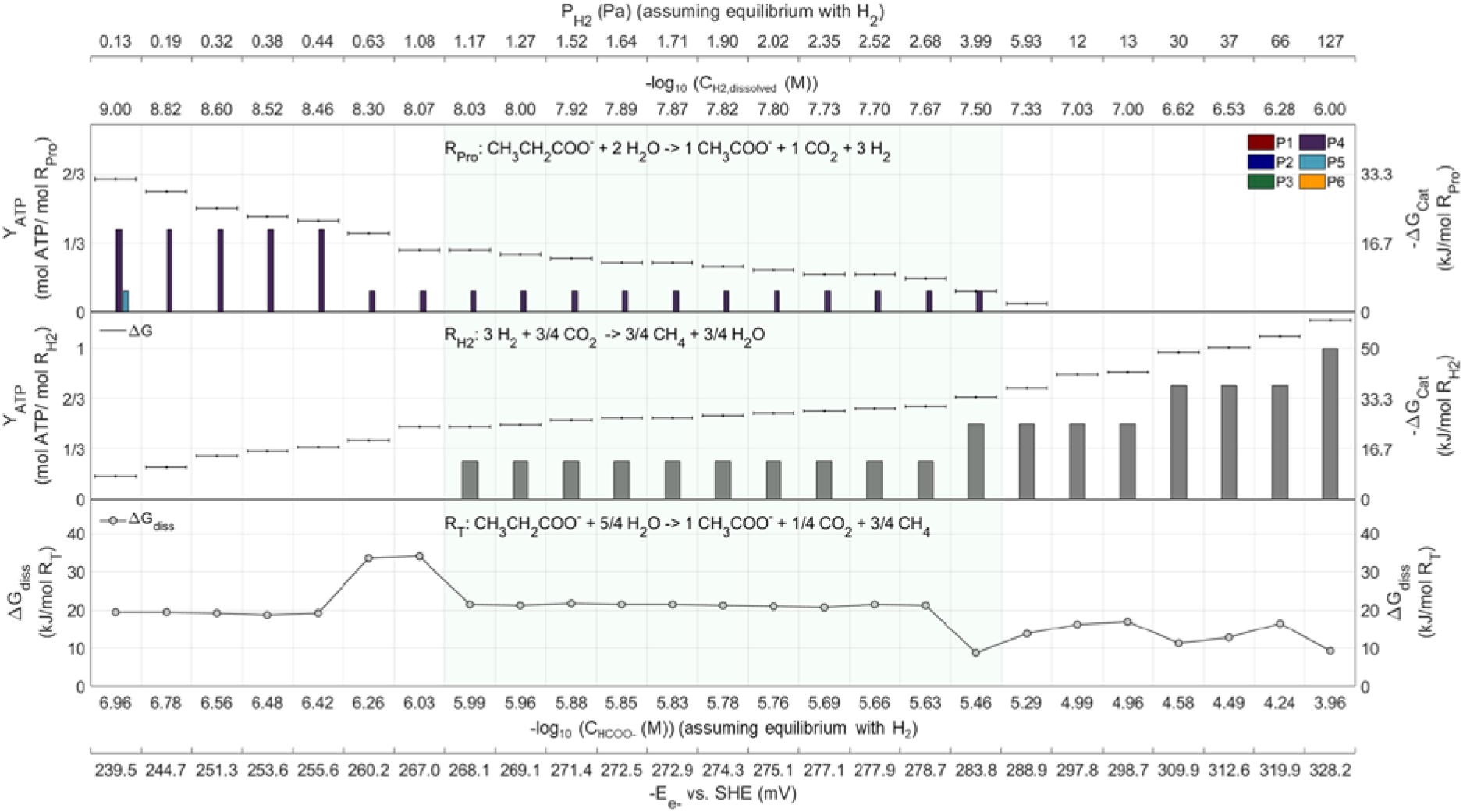
ATP yields for propionate oxidisers (top) and hydrogenotrophic methanogens (middle) as function of P_H2_ at 25 °C are shown in bars corresponding to feasible reactions with positive net ATP yields. Horizontal lines indicate the available catabolic energy. A range of hydrogen partial pressures is shown where both microbial functional groups can co-exist (shaded green area). The bottom plot shows the total energy dissipated (lost) in the complete syntrophic reaction. Alternative axis for IET via formate and direct electron transfer shows their values of concentration and voltage equivalent (in equilibrium) with the hydrogen concentrations and pressures shown.

The methanogenic niche appears to be limited to a range of P_H2_ (or equivalent alternative IET) between 1.2 and 4 Pa. This result based on the complete evaluation of all pathways variants for thermodynamic and physiological feasibility indicates that propionate oxidisers can only produce ATP within this hydrogen pressure range through the lactate pathway. Moreover, the region of highest overall reaction efficiency is found at P_H2_ between 2.7 and 4 Pa, coinciding with a leap up to higher ATP yield for the methanogen partner almost at the limit of feasibility for the propionate oxidiser.

Such low values of P_H2_ for syntrophic reaction feasibility are highly problematic as they are below the defined minimum physiological limit of 1 μM. Considering a bacterial cell volume of around 1 μm^3^, the number of hydrogen molecules present inside a cell within the methanogenic niche would be around 6 to 12. Such small number implies likely a kinetic impossibility for methanogenesis to actually take place. This suggests that IET between syntrophic partners should occur through alternative or additional mechanisms rather than through dissolved hydrogen. Moreover, if hydrogen is the only electron sink, the methylmalonyl-CoA pathway (the most studied pathway in propionate oxidation) does not appear to be feasible in any range of P_H2_ values evaluated. Sustained growth for the methanogen syntrophic partner, if based solely on dissolved hydrogen as electron donor, appears as an impossibility at such low concentrations as insufficient hydrogen is available.

The equivalent concentrations of formate, in equilibrium with hydrogen, calculated as alternative IET mechanism and shown in Figure 5 additional axis, show a more feasible methanogenic niche, since concentrations are above the defined lower physiological limit (1 μM). Alternatively, electrons could be transferred directly via a conductive material at potentials between −270 to −285 mV, as also shown in Figure 5.

## Conclusions

The results presented in this work indicate which pathway variations can provide the highest ATP yields for those microorganisms conducting propionate oxidation and hydrogenotrophic methanogenesis. The results are obtained based on a complete evaluation of the entire domain of possible pathway variations under constraints of known biochemistry, thermodynamic feasibility of all reactions steps and physiologically acceptable concentrations for all metabolites involved. Propionate oxidation via lactate pathways appears to have the highest ATP yields per mole of propionate. Our results suggest as well that the energy recovered through the methylmalonyl-CoA pathways does not suffice to sustain microbial life even in a methanogenic environment (P_H2_ = 1.62 Pa) under the conditions evaluated. Propionate oxidation only appears feasible via the lactate pathway when P_H2_ values are within the methanogenic niche. The extremely low P_H2_ values (below the minimum reasonable physiological limits) calculated for compatible simultaneous microbial growth of both syntrophs indicate that the IET via dissolved hydrogen is very unlikely, due to the extremely low number of hydrogen molecules found within the volume of a cell at those concentrations. These results suggest that IET must occur through alternative mechanisms such as via formate or via direct electron transfer (e.g. via conductive pili) at a voltage near −275 mV vs. SHE. The algorithm developed appears to be well suited for the study of other energy limited microbial metabolisms as it is largely founded on first principles and an entirely mechanistic basis.

## Acknowledgements

This publication is based upon work supported by Khalifa University’s Award No. CIRA-2018-84 and the Government of Abu Dhabi.

## References

1. Thauer RK, Jungermann K, Decker K. Energy conservation in chemotrophic anaerobic bacteria. Bacteriological reviews. 1977 Mar;41(1):100–80.

2. Weiss DS, Thauer RK. Methanogenesis and the unity of biochemistry. Cell. 1993 Mar 26;72(6):819–22.

3. Thauer RK, Kaster A-K, Seedorf H, Buckel W, Hedderich R. Methanogenic archaea: ecologically relevant differences in energy conservation. Nature Reviews Microbiology. 2008 30/online;6:579.

4. Stams AJ, Plugge CM. Electron transfer in syntrophic communities of anaerobic bacteria and archaea. Nature reviews Microbiology. 2009 Aug;7(8):568–77.

5. Jackson BE, McInerney MJ. Anaerobic microbial metabolism can proceed close to thermodynamic limits. Nature. 2002 Jan;415(6870):454–6.

6. Schink B. Energetics of syntrophic cooperation in methanogenic degradation. Microbiology and Molecular Biology Reviews. 1997;61(2):262–80.

7. Thauer RK. Biochemistry of methanogenesis: a tribute to Marjory Stephenson:1998 Marjory Stephenson Prize Lecture. Microbiology. 1998;144(9):2377–406.

8. Schmidt JE, Ahring BK. Effects of hydrogen and formate on the degradation of propionate and butyrate in thermophilic granules from an upflow anaerobic sludge blanket reactor. Appl Environ Microbiol. 1993 Aug;59(8):2546–51.

9. Boone DR, Johnson RL, Liu Y. Diffusion of the Interspecies Electron Carriers H(2) and Formate in Methanogenic Ecosystems and Its Implications in the Measurement of K(m) for H(2) or Formate Uptake. Applied and environmental microbiology. 1989;55(7):1735–41.

10. de Bok FAM, Plugge CM, Stams AJM. Interspecies electron transfer in methanogenic propionate degrading consortia. Water Research. 2004;38(6):1368–75.

11. Müller N, Worm P, Schink B, Stams AJM, Plugge CM. Syntrophic butyrate and propionate oxidation processes: from genomes to reaction mechanisms. Environmental Microbiology Reports. 2010;2(4):489–99.

12. Storck T, Virdis B, Batstone DJ. Modelling extracellular limitations for mediated versus direct interspecies electron transfer. The ISME Journal. 2016;10:621.

13. Schink B, Montag D, Keller A, Müller N. Hydrogen or formate: Alternative key players in methanogenic degradation. Environmental Microbiology Reports. 2017;9(3):189–202.

14. Dong X, Stams AJM. Evidence for H2 and formate formation during syntrophic butyrate and propionate degradation. Anaerobe. 1995 Feb 1;1(1):35–9.

15. Stams AJM, Dong X. Role of formate and hydrogen in the degradation of propionate and butyrate by defined suspended cocultures of acetogenic and methanogenic bacteria. Antonie van Leeuwenhoek. 1995;68(4):281–4.

16. Kosaka T, Kato S, Shimoyama T, Ishii S, Abe T, Watanabe K. The genome of Pelotomaculum thermopropionicum reveals niche-associated evolution in anaerobic microbiota. Genome Research. 2008 Mar 1;18(3):442–8.

17. Reguera G, McCarthy KD, Mehta T, Nicoll JS, Tuominen MT, Lovley DR. Extracellular electron transfer via microbial nanowires. Nature. 2005 Jun;435(7045):1098–101.

18. Shrestha PM, Rotaru A-E, Aklujkar M, Liu F, Shrestha M, Summers ZM, et al. Syntrophic growth with direct interspecies electron transfer as the primary mechanism for energy exchange. Environmental Microbiology Reports. 2013;5(6):904–10.

19. Rotaru A-E, Shrestha PM, Liu F, Shrestha M, Shrestha D, Embree M, et al. A new model for electron flow during anaerobic digestion: direct interspecies electron transfer to Methanosaeta for the reduction of carbon dioxide to methane. Energy Environ Sci. 2014;7(1):408–15.

20. Reguera G. Microbial nanowires and electroactive biofilms. FEMS Microbiol Ecol [Internet]. 2018 Jul 1 [cited 2019 Mar 6];94(7). Available from: https://academic.oup.com/femsec/article/94/7/fiy086/5000162

21. Morita M, Malvankar NS, Franks AE, Summers ZM, Giloteaux L, Rotaru AE, et al. Potential for Direct Interspecies Electron Transfer in Methanogenic Wastewater Digester Aggregates. mBio [Internet]. 2011 Sep 1 [cited 2020 Feb 6];2(4). Available from: https://mbio.asm.org/content/2/4/e00159-11

22. Kato S, Hashimoto K, Watanabe K. Microbial interspecies electron transfer via electric currents through conductive minerals. PNAS. 2012 Jun 19;109(25):10042–6.

23. Shrestha PM, Rotaru A-E. Plugging in or going wireless: strategies for interspecies electron transfer. Front Microbiol [Internet]. 2014 [cited 2019 Mar 7];5. Available from: https://www.frontiersin.org/articles/10.3389/fmicb.2014.00237/full

24. Lovley DR. Syntrophy Goes Electric: Direct Interspecies Electron Transfer. Annual Review of Microbiology. 2017;71(1):643–64.

25. Yamada C, Kato S, Ueno Y, Ishii M, Igarashi Y. Conductive iron oxides accelerate thermophilic methanogenesis from acetate and propionate. Journal of Bioscience and Bioengineering. 2015 Jun 1;119(6):678–82.

26. Koch M, Dolfing J, Wuhrmann K, Zehnder AJB. Pathways of Propionate Degradation by Enriched Methanogenic Cultures. Applied and environmental microbiology. 1983;45(4):1411–4.

27. Stams AJM, De Bok FAM, Plugge CM, Van Eekert MHA, Dolfing J, Schraa G. Exocellular electron transfer in anaerobic microbial communities. Environmental Microbiology. 2006;8(3):371–82.

28. Kosaka T, Uchiyama T, Ishii S, Enoki M, Imachi H, Kamagata Y, et al. Reconstruction and Regulation of the Central Catabolic Pathway in the Thermophilic Propionate-Oxidizing Syntroph *Pelotomaculum thermopropionicum*. Journal of Bacteriology. 2006;188(1):202–10.

29. Hamilton JJ, Calixto Contreras M, Reed JL. Thermodynamics and H2 Transfer in a Methanogenic, Syntrophic Community. PLoS Computational Biology. 2015;11(7):e1004364.

30. Hidalgo-Ahumada CAP, Nobu MK, Narihiro T, Tamaki H, Liu W-T, Kamagata Y, et al. Novel energy conservation strategies and behaviour of *Pelotomaculum schinkii* driving syntrophic propionate catabolism. Environmental Microbiology. 2018;0(0).

31. Kaziro Y, Ochoa S. The Metabolism of Propionic Acid. In: Advances in Enzymology and Related Areas of Molecular Biology [Internet]. John Wiley & Sons, Ltd; 1964 [cited 2019 Jan 30]. p. 283–378. Available from: https://onlinelibrary.wiley.com/doi/abs/10.1002/9780470122716.ch7

32. Tholozan JL, Touzel JP, Samain E, Grivet JP, Prensier G, Albagnac G. Clostridium neopropionicum sp. nov., a strict anaerobic bacterium fermenting ethanol to propionate through acrylate pathway. Archives of Microbiology. 1992 Feb 1;157(3):249–57.

33. Tholozan JL, Samain E, Grivet JP, Moletta R, Dubourguier HC, Albagnac G. Reductive carboxylation of propionate to butyrate in methanogenic ecosystems. Appl Environ Microbiol. 1988 Feb;54(2):441–5.

34. de Bok FAM, Stams AJM, Dijkema C, Boone DR. Pathway of Propionate Oxidation by a Syntrophic Culture of *Smithella propionica* and *Methanospirillum hungatei*. Applied and environmental microbiology. 2001;67(4):1800–4.

35. Kleerebezem R, Stams AJ. Kinetics of syntrophic cultures: a theoretical treatise on butyrate fermentation. Biotechnol Bioeng. 2000 Mar 5;67(5):529–43.

36. Rodríguez J, Kleerebezem R, Lema JM, van Loosdrecht MC. Modeling product formation in anaerobic mixed culture fermentations. Biotechnology and Bioengineering. 2006 Feb 20;93(3):592–606.

37. Bar-Even A, Flamholz A, Noor E, Milo R. Thermodynamic constraints shape the structure of carbon fixation pathways. Biochimica et Biophysica Acta (BBA) - Bioenergetics. 2012 Sep 1;1817(9):1646–59.

38. González-Cabaleiro R, Lema JM, Rodríguez J, Kleerebezem R. Linking thermodynamics and kinetics to assess pathway reversibility in anaerobic bioprocesses. Energy & Environmental Science. 2013;6(12):3780.

39. Noor E, Bar-Even A, Flamholz A, Reznik E, Liebermeister W, Milo R. Pathway Thermodynamics Highlights Kinetic Obstacles in Central Metabolism. PLoS Computational Biology. 2014 20 06/18/received 01/08/accepted;10(2):e1003483.

40. Regueira A, González-Cabaleiro R, Ofiţeru ID, Rodríguez J, Lema JM. Electron bifurcation mechanism and homoacetogenesis explain products yields in mixed culture anaerobic fermentations. Water Research. 2018 Sep 15;141:349–56.

41. Sieber JR, Sims DR, Han C, Kim E, Lykidis A, Lapidus AL, et al. The genome of Syntrophomonas wolfei: new insights into syntrophic metabolism and biohydrogen production. Environmental Microbiology. 2010;12(8):2289–301.

42. McInerney MJ, Sieber JR, Gunsalus RP. Microbial Syntrophy: Ecosystem-Level Biochemical Cooperation. Microbe Magazine. 2011 Jan 1;6(11):479–85.

43. Losey NA, Mus F, Peters JW, Le HM, McInerney MJ. Syntrophomonas wolfei Uses an NADH-Dependent, Ferredoxin-Independent [FeFe]-Hydrogenase To Reoxidize NADH. Appl Environ Microbiol. 2017 Oct 15;83(20):e01335–17.

44. Batstone DJ, Picioreanu C, van Loosdrecht MCM. Multidimensional modelling to investigate interspecies hydrogen transfer in anaerobic biofilms. Water Research. 2006;40(16):3099–108.

45. Rouvière PE, Wolfe RS. Novel biochemistry of methanogenesis. J Biol Chem. 1988 Jun 15;263(17):7913–6.

46. Wolfe RS. My Kind of Biology. Annual Review of Microbiology. 1991;45(1):1–36.

47. Thauer RK. The Wolfe cycle comes full circle. Proceedings of the National Academy of Sciences. 2012;109(38):15084–5.

48. Becher B, Müller V, Gottschalk G. N5-methyl-tetrahydromethanopterin:coenzyme M methyltransferase of Methanosarcina strain Gö1 is an Na(+)-translocating membrane protein. Journal of Bacteriology. 1992;174(23):7656–60.

49. Kaster A-K, Moll J, Parey K, Thauer RK. Coupling of ferredoxin and heterodisulfide reduction via electron bifurcation in hydrogenotrophic methanogenic archaea. PNAS. 2011 Feb 15;108(7):2981–6.

50. Kanehisa M, Goto S. KEGG: kyoto encyclopedia of genes and genomes. Nucleic acids research. 2000 Jan 1;28(1):27–30.

51. Rosen C, Jeppsson U. Aspects on ADM1 Implementation within the BSM2 Framework. Department of Industrial Electrical Engineering and Automation, Lund University, Lund, Sweden. 2006;1–35.

52. Bennett BD, Kimball EH, Gao M, Osterhout R, Van Dien SJ, Rabinowitz JD. Absolute metabolite concentrations and implied enzyme active site occupancy in Escherichia coli. Nature chemical biology. 2009 Aug;5(8):593–9.

53. Atkinson D. Cellular energy metabolism and its regulation. New York: Academic Press; 1977. xi, 293 p.

54. Milo R. What is the total number of protein molecules per cell volume? A call to rethink some published values. BioEssays : news and reviews in molecular, cellular and developmental biology. 2013;35(12):1050–5.

55. Reich J, Selkov E. Energy Metabolism of the Cell. A Theoretical Treatise. London: Academic Press; 1981.

56. Hanselmann KW. Microbial energetics applied to waste repositories. Experientia. 1991;47(7):645–87.

57. Stephanopoulos G, Aristidou AA, Nielsen JH. Metabolic engineering : principles and methodologies. San Diego: Academic Press; 1998. xxi, 725 p.

58. Alberty RA. Biochemical thermodynamics: applications of Mathematica. Methods of biochemical analysis. 2006;48:1–458.

59. Flamholz A, Noor E, Bar-Even A, Milo R. eQuilibrator—the biochemical thermodynamics calculator. Nucleic acids research. 2012;40(D1):D770–5.

60. Dimroth P. The Role of Biotin and Sodium in the Decarboxylation of Oxaloacetate by the Membrane-Bound Oxaloacetate Decarboxylase from Klebsiella aerogenes. European Journal of Biochemistry. 1982;121(2):435–41.

61. Hilpert W, Dimroth P. Conversion of the chemical energy of methylmalonyl-CoA decarboxylation into a Na+gradient. Nature. 1982 08/online;296:584.

62. Gottschalk G, Thauer RK. The Na+-translocating methyltransferase complex from methanogenic archaea. Biochimica et Biophysica Acta (BBA) - Bioenergetics. 2001 May 1;1505(1):28–36.

63. Hilpert W, Schink B, Dimroth P. Life by a new decarboxylation-dependent energy conservation mechanism with Na as coupling ion. The EMBO journal. 1984;3(8):1665–70.

64. Buckel W, Thauer RK. Flavin-Based Electron Bifurcation, Ferredoxin, Flavodoxin, and Anaerobic Respiration With Protons (Ech) or NAD+ (Rnf) as Electron Acceptors: A Historical Review. Frontiers in Microbiology. 2018 Mar;9(401).

65. Müller V, Chowdhury NP, Basen M. Electron Bifurcation: A Long-Hidden Energy-Coupling Mechanism. Annu Rev Microbiol. 2018 Sep 8;72(1):331–53.

66. Caspi R, Billington R, Ferrer L, Foerster H, Fulcher CA, Keseler IM, et al. The MetaCyc database of metabolic pathways and enzymes and the BioCyc collection of pathway/genome databases. Nucleic Acids Res. 2016 Jan 4;44(D1):D471–80.

67. Buckel W, Thauer RK. Flavin-Based Electron Bifurcation, A New Mechanism of Biological Energy Coupling. Chem Rev. 2018 Apr 11;118(7):3862–86.

68. Nakanishi-Matsui Mayumi, Futai M. Stochastic rotational catalysis of proton pumping F-ATPase. Philosophical Transactions of the Royal Society B: Biological Sciences. 2008 Jun 27;363(1500):2135–42.

69. Cross RL, Müller V. The evolution of A-, F-, and V-type ATP synthases and ATPases: reversals in function and changes in the H+/ATP coupling ratio. FEBS Letters. 2004 Oct 8;576(1):1–4.

70. Mitome N, Suzuki T, Hayashi S, Yoshida M. Thermophilic ATP synthase has a decamer c-ring: Indication of noninteger 10:3 H+/ATP ratio and permissive elastic coupling. Proc Natl Acad Sci U S A. 2004 Aug 17;101(33):12159–64.

71. Pogoryelov D, Klyszejko AL, Krasnoselska GO, Heller E-M, Leone V, Langer JD, et al. Engineering rotor ring stoichiometries in the ATP synthase. PNAS. 2012 Jun 19;109(25):E1599–608.

72. Mayer F, Müller V. Adaptations of anaerobic archaea to life under extreme energy limitation. FEMS Microbiol Rev. 2014 May 1;38(3):449–72.

73. Lever MA, Rogers KL, Lloyd KG, Overmann J, Schink B, Thauer RK, et al. Life under extreme energy limitation: a synthesis of laboratory- and field-based investigations. FEMS Microbiol Rev. 2015 Sep 1;39(5):688–728.

74. Hoehler TM, Alperin MJ, Albert DB, Martens CS. Apparent minimum free energy requirements for methanogenic Archaea and sulfate-reducing bacteria in an anoxic marine sediment. FEMS Microbiology Ecology. 2001 Dec 1;38(1):33–41.

75. Boynton ZL, Bennett GN, Rudolph FB. Intracellular Concentrations of Coenzyme A and Its Derivatives from Clostridium acetobutylicum ATCC 824 and Their Roles in Enzyme Regulation. Appl Environ Microbiol. 1994 Jan 1;60(1):39–44.

